# Transcriptional burst kinetics are linked to short term transcriptional memory

**DOI:** 10.1101/2021.10.31.466715

**Authors:** Adrien Senecal, Robert Singer, Robert Coleman

## Abstract

Transcriptional bursting is thought to be a stochastic process that allows the dynamic regulation of most genes. The random telegraph model assumes the existence of two states, ON and OFF. However recent studies indicate the presence of additional ON states, suggesting that bursting kinetics and their regulation can be quite complex. We have developed a system to study transcriptional bursting in the context of p53 biology using the endogenous *p21* gene tagged with MS2 in human cells. Remarkably, we find that transcriptional bursts from the *p21* gene contain multiple ON and OFF states that can be regulated by elevation of p53 levels. Distinct ON states are characterized by differences in burst duration, classified as Short and Long, with long bursts associated with higher Pol II initiation rates. Importantly, the different ON states display memory effects that allow us to predict the likelihood of properties of future bursting events. Long bursting events result in faster re-activation, longer subsequent bursts and higher transcriptional output in the future compared to short bursts. Bursting memory persists up to 2 hours suggesting a stable inheritable promoter architecture. Bursting memory at the *p21* gene is the strongest under basal conditions and is suppressed by UV and inhibition of H3K9me1/2, which also increase transcriptional noise. Stabilization of p53 by Nutlin-3a partially reverses suppression of bursting memory suggesting that higher p53 levels may be a key in enforcing memory under conditions of cellular stress. Overall our data uncover a new found bursting property termed Short-Term Transcriptional Memory (STTM) that has the potential to fine-tune transcriptional output at the *p21* gene.

## INTRODUCTION

Transcriptional bursting is often described as a stochastic process whereby a gene rapidly cycles between transcriptionally active and inactive states(Rodriguez and Larson 2020; Tunnacliffe and Chubb 2020). This stochastic nature of bursting is thought to generate transcriptional noise or heterogeneity in a population of cells(Raj and van Oudenaarden 2008; Eldar and Elowitz 2010). Regulation of transcriptional bursting and noise can be studied via many assays including smFISH and scRNA-seq(Patange et al. 2018; Larsson et al. 2019). However, examining transcription bursting in a live cell allows the measurement of temporal switching of a single loci between transcriptionally active and inactive states. Therefore, dynamic live single-cell transcriptional bursting measurements provide information that can’t readily be achieved by SNAP-shot assays in single cells (e.g. smFISH/scRNA-seq) or bulk populations (RT-qPCR)(Lenstra et al. 2016).

The random telegraph model of transcriptional bursting assumes the existence two states, ON and OFF(Lenstra et al. 2016). This model likely underestimates the level of complexity and regulation that exists when nearly a hundred proteins dynamically assemble and disassemble on a promoter and terminator to initiate or end a transcriptional burst(Biswas et al. 2021). In addition, the presence of alternative transcription pre-initiation scaffolds (e.g TFIID and SAGA directed) and core-promoter variants (e.g TATA and TATA-less) likely suggests additional layers of complexity(Huisinga and Pugh 2004; de Jonge et al. 2017; Kubik et al. 2017; Donczew et al. 2020; Timmers 2021). Indeed, recent live-cell studies revealed the presence of additional ON states during transcription of the β-actin gene indicating that burst kinetics are rather complex(Corrigan et al. 2016). Correspondingly, this affords the cell many different access points or mechanisms to dynamically regulate transcriptional output of a gene.

Numerous studies have shown that transcriptional output can be regulated by changes in burst duration (ON state), frequency (OFF state), and Pol II initiation rates (burst intensity) (Figure 1A)(Molina et al. 2013; Corrigan and Chubb 2014; Senecal et al. 2014). Transcriptional activators have been shown to increase burst duration, frequency and Pol II initiation rates. This makes sense since transcriptional activators recruit many factors (e.g. chromatin remodelers, TFIID, Mediator, Pol II, etc.) that are involved in turning on and off transcription. However, very few of these studies examined dynamic activator induced changes in an endogenous gene in the context of multiple ON state models. Therefore studying activator induced changes in multiple ON states within the context of precisely timed triggers on an endogenous gene could reveal detailed insights into the mechanistic regulation of bursting.

**Figure 1.**
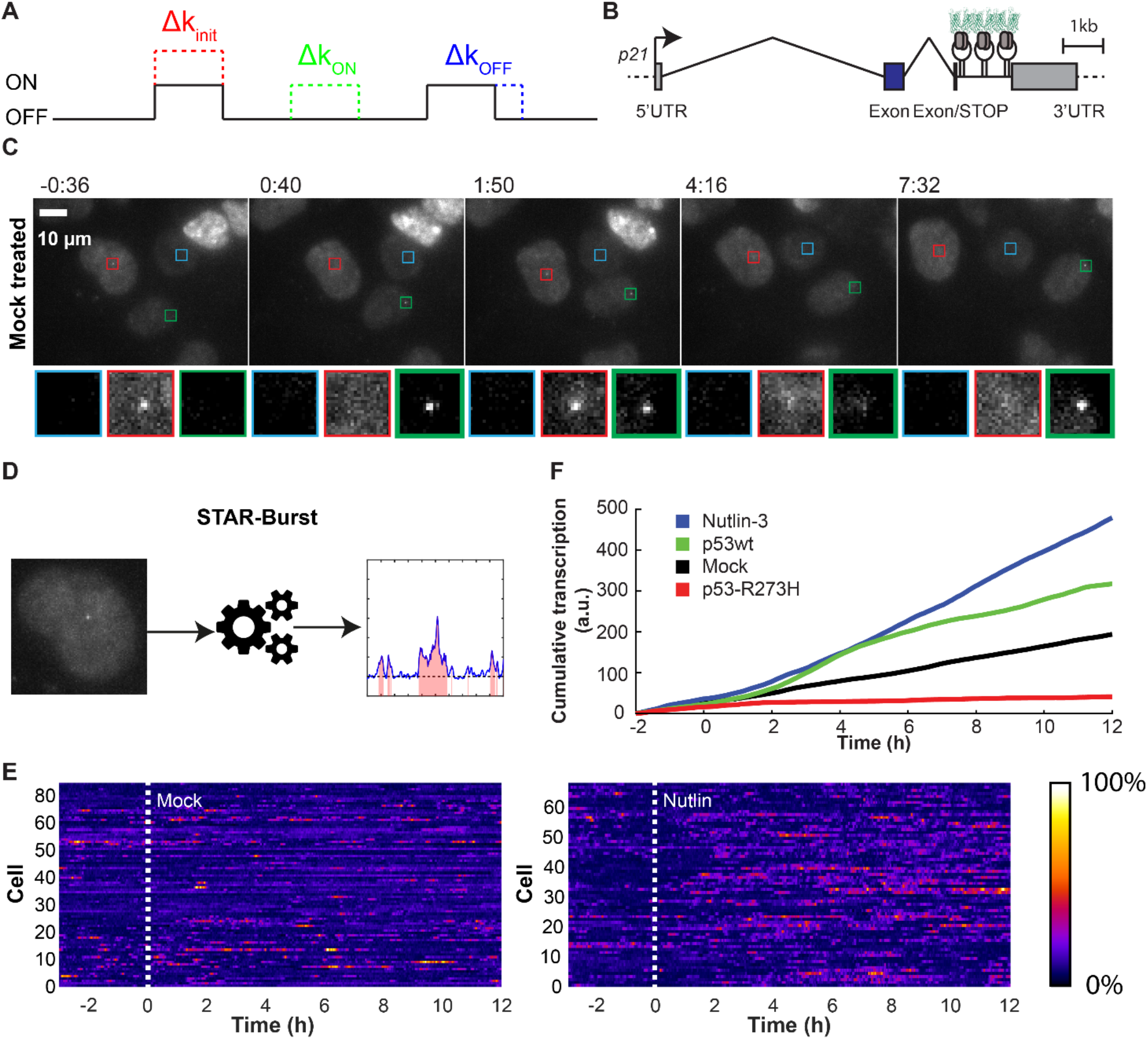
Live cell quantification of the endogenous *p21* expression. **(A)** Schematic representation of the random telegraph model for transcriptional bursting. Gene can switch between inactive (OFF) and active (ON) state. Transitions are described by rate constants k_on_ and k_off_. Transcripts are produced during ON states as a Poisson process with fixed rate, k_init_. **(B)** Schematic representation of the *p21*-MS2 gene. Transcription start site is shown as an arrow, exons are shown as rectangle, DNA coding regions are in blue while 5’ and 3’ UTR are in gray, and introns are shown as broken lines. 24 repeats of MS2 binding site RNA stem-loop (MS2-SL) have been inserted by CRISPR/Cas9 in 3^rd^ exons, just after the stop codon. GFP-tagged MS2 Capsid Protein (MCP-GFP) binds MS2-SL to visualize nascent transcription sites (TS). The scale represent 1000 base (1kb). **(C)** Live-cell imaging of U2OS *p21-MS2* cells. Images of the cell line with MS2 inserted into a single allele of the *p21* gene (maximum intensity projections of z-dimension stacks). The scale bar represent 10 μm. **(D)** Schematic representation of the STAR-Burst program. **(E)** TS intensities over time. Each lane is a cell and the TS intensity is color coded (colormap fire). Cells are treated after 3h with a mock solution (left panel) or Nutlin-3a (right panel). **(F)** Cumulative transcription of *p21-MS2.* The average cumulative transcription (above background) shows that Nutlin-3a (blue line, n = 68) and cells over-expressing p53 (green line, n = 94) transcribe the *p21* gene faster than untreated cells (black line, n = 85). The over-expression of p53-R273H exhibit a dominant-negative activity over the wild-type p53 (red line, n = 25).

The p53 mediated gene expression network is an ideal system for studying dynamic changes in gene expression since it can be activated by a number of stimuli and cellular stresses. Treatment of cells with different DNA damaging agents (e.g. UV or Doxorubicin) results in a mixture of responses ranging from cell cycle arrest via expression of the *p21* gene or apoptosis(Donner et al. 2007). Addition of Nutlin-3a to cells inhibits mdm2-mediated degradation of p53 which in turn causes elevation of p53 levels by non-genotoxic means to exclusively turn on cell cycle arrest via *p21* expression. Recent high-resolution genome-wide studies of p53 binding has shown stimuli-specific occupancy of p53 to its Response Elements (REs) to regulate overlapping and distinct sets of gene targets(Chang et al. 2014). Upon exposure to different stimuli, p53 can even bind to different REs controlling expression of the same gene, including the *p21* gene(Chang et al. 2014). Therefore various stimuli may be able to differentially regulate or fine tune p53-target gene expression via multiple mechanisms. This creates an opportunity to study how different stimuli affect bursting patterns of multiple ON states in a live cell.

In this paper, we establish a system for monitoring the dynamic expression of the *p21* gene under basal conditions and when activated via different stimuli. We first tested the system by examining how bursting profiles change at low and high p53 levels using a two state, ON/OFF, random telegraph model. With this two state model, elevation of p53 levels leads to increased burst duration, frequency and transcriptional output. By carefully analyzing burst duration and frequency, we determined that bursting of the *p21* gene displays multiple ON and OFF states. Further analysis of the different ON states revealed that bursts longer than 16 minutes had increased Pol II initiation rates compared to short bursts. Bursting patterns of the different ON states revealed properties of memory where bursts re-initiated faster and for longer periods of time after an initial long burst compared with after a short burst. Importantly, treating the cells with different stimuli (UV and inhibition of EHMT1/2) suppressed memory and increased transcriptional noise compared to basal conditions. Elevation of p53 levels during inhibition of EHMT1/2, which dimethylates H3K9 and p53, partially restored transcriptional bursting memory suggesting that increasing p53 levels may be a mechanism to buffer against memory perturbations.

## RESULTS

### Establishment of a system for quantification of dynamic inducible endogenous gene expression to study p53 biology in a living cell

To observe transcriptional regulation of endogenous alleles in single cells, we used the MS2 system which exploits a highly specific binding of MS2 capsid protein (MCP) to the MS2 stem loop (MS2-SL) (Vera et al., 2016). Using CRISPR-Cas9, we previously integrated 24 copies of the MS2-SL into the 3’ untranslated region (3’UTR) of CDKN1a gene in U2OS cells (*p21* - **Figure 1B**)(Carvajal et al. 2018). We validated the precise integration of MS2-SL repeats into the *p21* gene by PCR and smFISH(Carvajal et al. 2018). This genetically engineered cell line also stably expressed a green fluorescent protein (GFP)–tagged MS2 capsid protein that specifically recognizes MS2-SL tagged *p21* mRNA (MCP-GFP – **Figure 1B**). This system allowed us to directly measure transcriptional bursting dynamics from a canonical p53 target gene locus within its native genomic context in response to p53 activation (**Figure 1A**).

Live-cell imaging indicates that the *p21* mRNA is transcribed dynamically over short periods of time, defined as a series of transcriptional bursts (**Video and Figure 1C**). To quantify this cycling between active/inactive transcription, we tracked transcription sites (TSs) and measured the fluorescence intensity over time. We developed a program, STAR-burst (Site of Transcription, Analysis and Recapitulation of Bursts), to semi-automatically detect the TSs and quantify their activity over time (**Figure 1D and Figure S1**). Briefly, STAR-burst uses the images from the microscope, filters them to remove background noise, and tracks the TS looking for the maximum intensity pixel in a restricted area. The traces obtained are normalized based on periods of inactivity. The allele is assumed to be active if the fluorescence signal is greater than 1.5 fold above the normalized background.

At low p53 protein levels, individual cells show sporadic expression activity of the *p21* gene, with brief expression periods surrounded by much longer periods of inactivity (**Figure 1E**). Using Nutlin-3a, which increases the endogenous p53 protein concentration by inhibiting p53’s degradation by mdm2, we noticed an increase in the number and duration of transcription bursts (**Figure 1E – left panel**). Increased transcriptional activity of *p21* and production of more mRNA was seen when p53 levels were also elevated by exogenous over-expression of a p53 expressing transgene (**Figure 1F**). This suggests that the p53 protein is positively regulating transcriptional bursting from the *p21* gene. Interestingly, the transcriptional activity of *p21* is completely inhibited with virtually no *p21* mRNA produced when an oncogenic p53 mutant (p53-R273H) is overexpressed (**Figure 1F**). These results are consistent with previous studies showing that p53-R273H may act in a dominant negative manner since the mutant protein lacks the ability to recognize p53 Response Elements (p53 REs) in vitro yet still retains the ability to tetramerize with endogenous wild type p53(Bullock et al. 2000; Willis et al. 2004).

### Live-cell imaging of endogenous *p21* transcription reveals that p53 controls transcription burst frequency and duration

Using a random telegraph two state model of ON and OFF states, live single cell imaging reveals that *p21* is expressed with short burst of transcription followed by longer OFF period at low endogenous p53 protein levels (**Figure 2A**). After p53 protein stabilization with Nutlin-3a treatment, we observed an increase in the number (i.e. frequency) and duration of transcriptional bursts (**Figure 2A**). To quantify the induction of *p21* transcriptional bursting with respect to p53 protein levels, we measured the probability of *p21* TSs to be active in single cells containing both low and high p53 protein levels. At low p53 protein levels under basal conditions, there is 15% chance for the *p21* TS to be ON (**Figure 2B**). The probability of a *p21* TS being active increases drastically to about 40% after both p53 over-expression and Nutlin-3a induction. Interestingly, the probability of a *p21* TS being active decreases after four hours of p53 over-expression. This can be explained by p53’s transcriptional induction of *mdm2*, which acts as an ubiquitin ligase and covalently attaches ubiquitin to p53 for degradation thus creating a negative feedback loop(Lev Bar-Or et al. 2000). Correspondingly, this negative feedback loop can be disrupted by treatment of cells with Nutlin-3a, which inhibits p53’s interaction with mdm2(Vassilev et al. 2004). In such cases, the probability of a *p21* TS being active remains high and stable for many hours after p53 stabilization with Nutlin-3a (**Figure 2B**).

**Figure 2.**
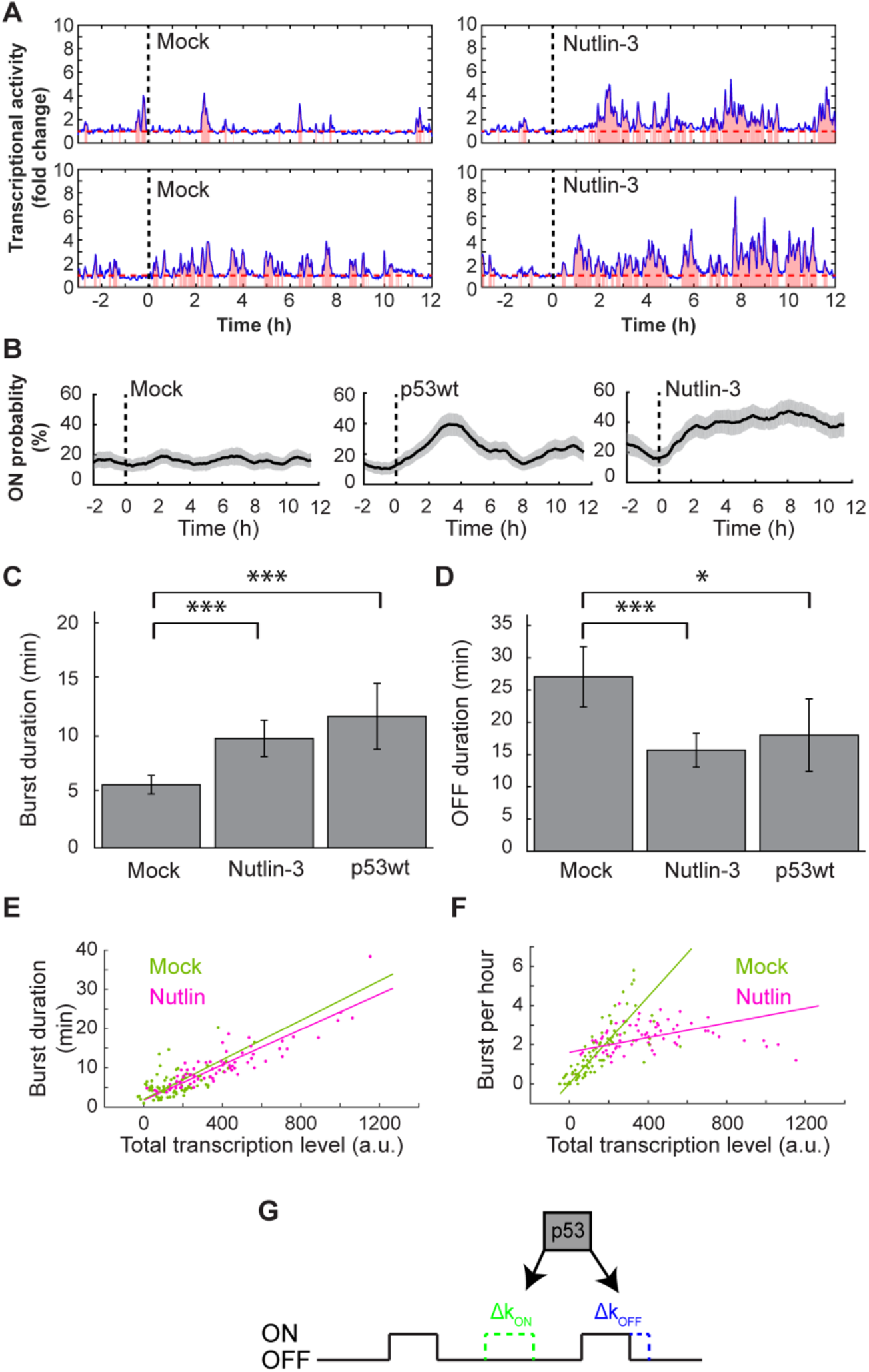
p53 increase p21 expression by enhancing burst duration and frequency. **(A)** Example of quantification of the fluorescence intensity of the TS treated with a mock solution or Nutlin-3a. The fluorescence intensity level is divided to the average background level to obtain a fold change. When the fluorescence intensity is higher than 1.5 fold the background, *p21* is deemed active (pink). **(B)** Proportion of cells containing an active TS (≥ 1.5 fold the background). For each time point and condition, the proportion of active TS calculated over a sliding window of 30 minutes. The grey area represents the 95% confidence interval. **(C and D**) Average burst duration and delay between bursts calculated for a 2h window (between 4 and 6 hours after induction). An increase of p53 concentration enhances burst duration and reduces delay between bursts (increase burst frequency). * for p < 0.05, ** for p-value < 0.01 and *** for p < 10^-3^ for all figures. Error bars represent the 95% confidence interval for all figures. **(C)** Average burst duration (ON duration). **(D)** Average duration of the interval between bursts (OFF duration). **(E)** Scatter plot of the burst duration in function of the total transcription level. The control cells are plotted in green and the Nutlin-3 treated-cells are plotted in fuchsia. Control and Nutlin-3a treated cells are fit with similar simple linear regression with respective r-values of 0.56 and 0.88). **(F)** Scatter plot of the burst frequency (burst per hours) as a function of the total transcription level. The control cells are plotted in green and the Nutlin-3 treated-cells are plotted in fuchsia. Control and Nutlin-3a treated cells are fit with a simple linear regression model with respective r-values of 0.75 and 0.16. **(G)** p53 controls both the burst frequency (k_ON_) and the burst duration (k_OFF_) of the *p21* gene.

To understand how the transcription of *p21* is dynamically regulated, we quantified the transcription burst duration (**Figure 2C**) and the delay between two bursts (OFF duration – **Figure 2D**). We observed that the *p21* transcription burst duration increases at high p53 protein levels (**Figure 2C**, 9.7 ± 1.6 min for Nutlin-3a and 11.7 ± 2.9 min for p53 over-expression versus 5.6 ± 0.8 min after Mock treatment). This data suggests that p53 increases transcriptional output from the *p21* gene by maintaining the promoter in an ON state for longer periods of times. The frequency of transcriptional bursting is reflected by the OFF duration between two bursts. Similarly, we found that the frequency of transcriptional bursting from the *p21* gene increases at high p53 protein levels (**Figure 2D**, OFF duration is 15.7 ± 2.6 min after Nutlin-3a and 18.0 ± 5.6 min after p53 over-expression versus 27.0 ± 4.7 min after mock treatment). Our live cell experiment allows us to follow changes in burst kinetics change over-time. Due to a likely negative feedback of MDM2 on p53 levels, burst duration and burst frequency are more transiently affected after p53 over-expression than Nutlin-3a induction (**Figures S2A and S2B**). These data suggest that p53 regulates transcriptional output of the *p21* gene by increasing the conversion of the promoter from an OFF to an ON state (i.e. frequency), while also extending the time that the promoter stays in an ON state (i.e. duration) (**Figure 2G**).

During a burst, the total transcriptional output is a function of the frequency of RNA Pol II initiation (e.g. Pol II flux) and the duration of the ON state. To determine if p53 affects Pol II flux, we compared the burst duration to the total transcription level for both mock (low p53) and Nutlin-3a treated (high p53) cells (**Figures 2E**). The burst duration at both low and high p53 protein levels was linearly correlated with the total number of mRNAs produced (r = 0.56 for Mock and r = 0.88 for Nutlin-3). Based on this linear correlation, there are no apparent differences in Pol II flux since the slopes are nearly equivalent at low and high p53 protein levels.

Our data also indicated that p53 increases the frequency of transcriptional bursting (**Figure 2D**). Therefore, we additionally examined if the frequency of bursts correlated with transcriptional output. While the frequency of the bursts is strongly correlated (r = 0.75) with the total transcription output at low p53 protein levels, this correlation disappears (r = 0.16) when p53 protein levels are high (**Figure 2F**). However at high p53 protein levels, the transcriptional output remains at a very high level which is much greater than the majority of bursts in untreated cells. The transcription frequency saturates around 2-3 bursts per hour at high p53 protein levels and cannot be increased beyond this with a simple increase of p53 concentration. This is likely due to the fact that p53 significantly extends the burst duration (**Figure 2C**) placing an upper limit on the number of burst cycles that are able to occur in a hour. Taken together, these results suggest a control of the overall *p21* transcription output by p53 by a regulation of the bursts duration and their frequency, up to a certain level (**Figure 2G**).

### Multiple ON and OFF states regulate *p21* transcriptional bursting

A previous study examining transcriptional bursting of the β-actin gene revealed the existence of multiple ON states during a transcriptional burst(Corrigan et al. 2016). To see if there were multiple ON states regulating *p21* transcription, we fit 1-CDF histograms of ON durations to models for a single or double exponential decay. Statistical analysis revealed two different classes of ON states with 85% of the bursts being short (4.3 ± 0.2 min) and 15% long (16.8 ± 1.9 min) under basal conditions **(Figure 3A)**. Additional fitting techniques using GRID analysis(Reisser et al. 2020) also indicated multiple populations of ON states **(Figure S3A)**. The most prevalent ON state found by GRID roughly matches the duration of the short burst found in multi-exponential fitting ( 2.8 ± 0.2 vs 4.3 ± 0.2 min). The percentage of events falling into longest duration ON state (~22 min) found by GRID analysis was poorly fit (4.8 ± 3.5%) after resampling. This indicates that multi-exponential fitting and GRID analysis find multiple ON states. However GRID analysis has indicated issues resolving the longer lived burst ON states likely due to the number of long burst events seen. Therefore to be conservative, ON states were defined using multiexponential fitting. Elevation of p53 levels via overexpression significantly increased the ratio of Long to Short bursts, without varying the duration of the two ON states indicating that p53 overexpression can lengthen a burst by altering the distribution of ON states **(Figures 3B and 3C)**.

**Figure 3.**
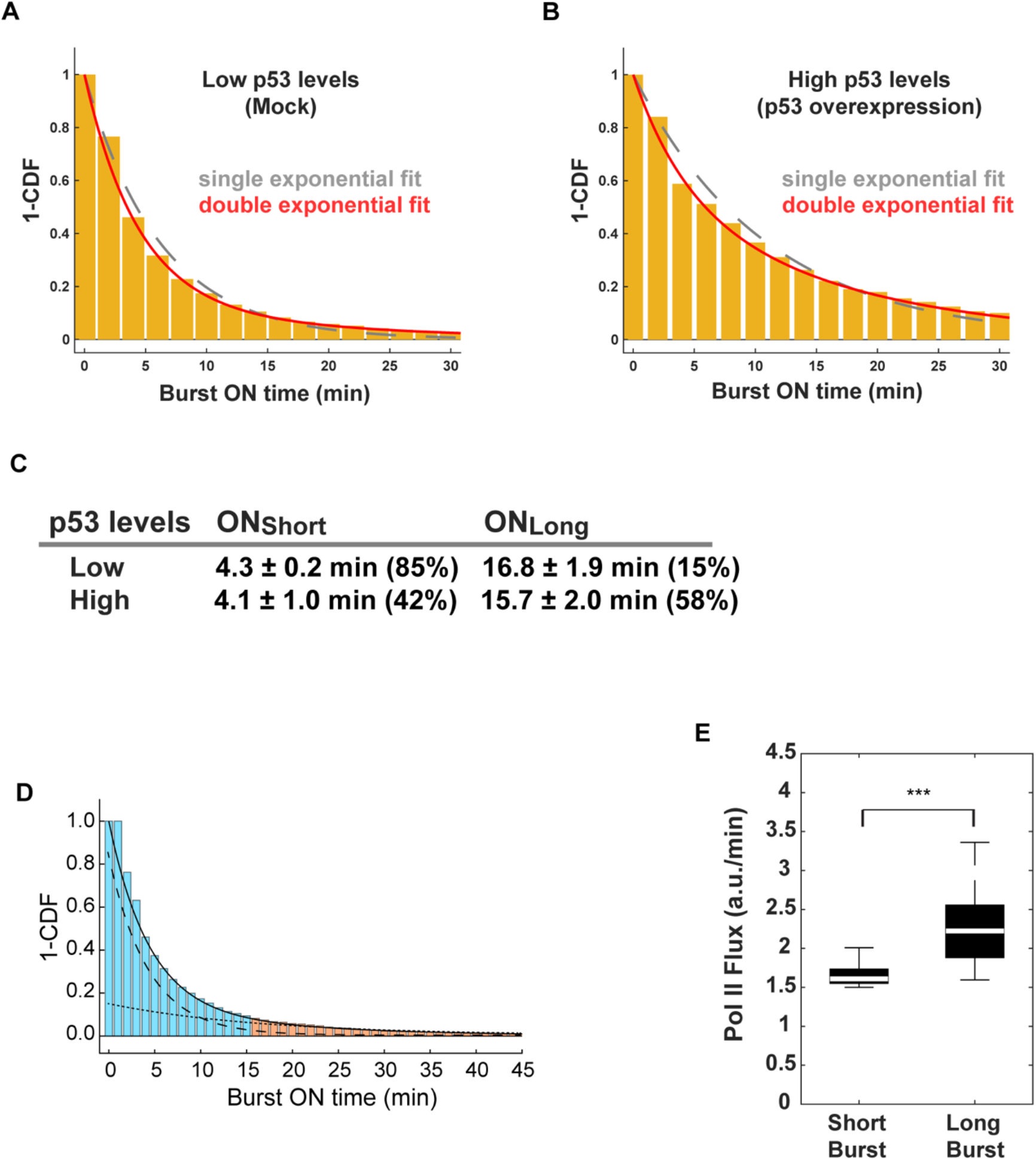
Bursting displays multiple ON states associate with different Pol II initiation rates. **(A-B)** ON duration can be split in two populations (short bursts and long bursts). Histogram (1-CDF) of the repartition of the ON duration during 15 hours of Mock treated cells **(A)** and 12 hours of p53 overexpression **(B)**. The single (dashed gray) and double exponential (the solid red) fits are shown. **(C)** Table showing the percentage of events and average duration of the short and long bursts for Mock treated cells and p53 overexpression. **(D)** For Mock treated cells, the sum of double exponential (the full line) are fitted to the ON duration: a short population (the long dashed line - 4.3 min 85% of events) and a long population (the short dashed line - 16.7 min 15% of events). Less than 2.5% of short event are longer than 15 min. We define short bursts (blue) as an ON duration of 15 min or less and long bursts (orange) as ON duration of more than 15 min. **(F)** Box plot of the Pol II flux (e.g. Pol II initiation rate) during a short and long burst.). p-value *** = < 0.001. The white bar in the solid black box is the median, while the lower and upper boundaries of the black box correspond to the 25^th^ and 75^th^ percentiles of the data respectively. Outliers typically represented less than 10% of the dataset and therefore were omitted for clarity.

To determine if there were multiple OFF states associated with *p21* transcriptional bursting, we performed GRID analysis of OFF durations since preliminary multi-exponential fitting revealed at least 3 populations of OFF states. GRID analysis revealed 4 different classes of OFF durations ranging from approximately three minutes to three hours under basal conditions **(Figure S3B)**. Elevation of p53 levels via overexpression eliminated the long-lived (165 min) OFF class and created two new well defined shorter (66 min and 6 min) OFF states compared to basal conditions. These results suggest that p53 may help to maintain the promoter in a semi-poised state for faster reactivation of a burst via an unknown mechanism. Overall, the presence of multiple ON and OFF states in transcriptional bursting profiles at the *p21* gene, which we term multi-phasic transcriptional bursting (MPTB), suggests much more complex regulatory mechanisms than seen in previous studies on transcriptional bursting.

### Pol II initiation rates differ between the two ON states

To characterize how transcriptional output differed between the short and long ON states, we compared the Pol II initiation rates (e.g. Pol II flux) associated with short and long bursts. Our previous multi-exponential fitting of ON duration histograms indicated that there was a ~4 fold difference in the length of the bursts in the different ON states. This large difference in durations amongst the different burst populations allowed us to classify individual burst events into either short or long bursts via thresholding. A 15 minute threshold was used to classify long burst events, since only 2.5% of short burst are longer that this duration under basal conditions **(Figure 3D)**. It is important to note that this threshold varies based upon the percentage of burst events in the short population and the duration of the short burst as determined via multi-exponential fitting methods (see methods). The cumulative transcriptional output from each individual short or long burst was then divided by the burst duration to yield a Pol II initiation rate (a.u./min). Median Pol II initiation rates for events classified as a short bursts were only slightly higher (1.6 a.u./min) than the threshold (1.5 fold above background) set to define transcription as being in an ON state of bursting **(Figure 3E)**. In contrast, the median Pol II initiation rates during a long burst were significantly higher (2.2 a.u./min) compared to short bursts suggesting that increased Pol II flux was associated with increased burst durations.

### Persistent short term memory of transcriptional activity

Many inducible genes display transcriptional memory that is typically measured after a stimulus over hours or after mitosis(D’Urso and Brickner 2017; Palozola et al. 2019). We next wanted to know if a particular ON state is persistent across many burst cycles. To see if TSs keep a memory of past transcriptional events during subsequent burst cycles, we decided to determine the probability for different sequences of bursts under basal conditions. A global analysis of the data indicates that two sequential long bursts were predominantly seen in our data compared to a short followed by a long burst (**Figures 4A, 17.4% vs 7.3%**). To quantitatively define this memory, we created a bursting memory index (BMI, observed/expected) associated with the two classes of burst durations. The memory index indicates that a train of long bursts (e.g. two sequential) is enriched in our data, while a long burst arising from a short burst is slightly disfavored.

**Figure 4.**
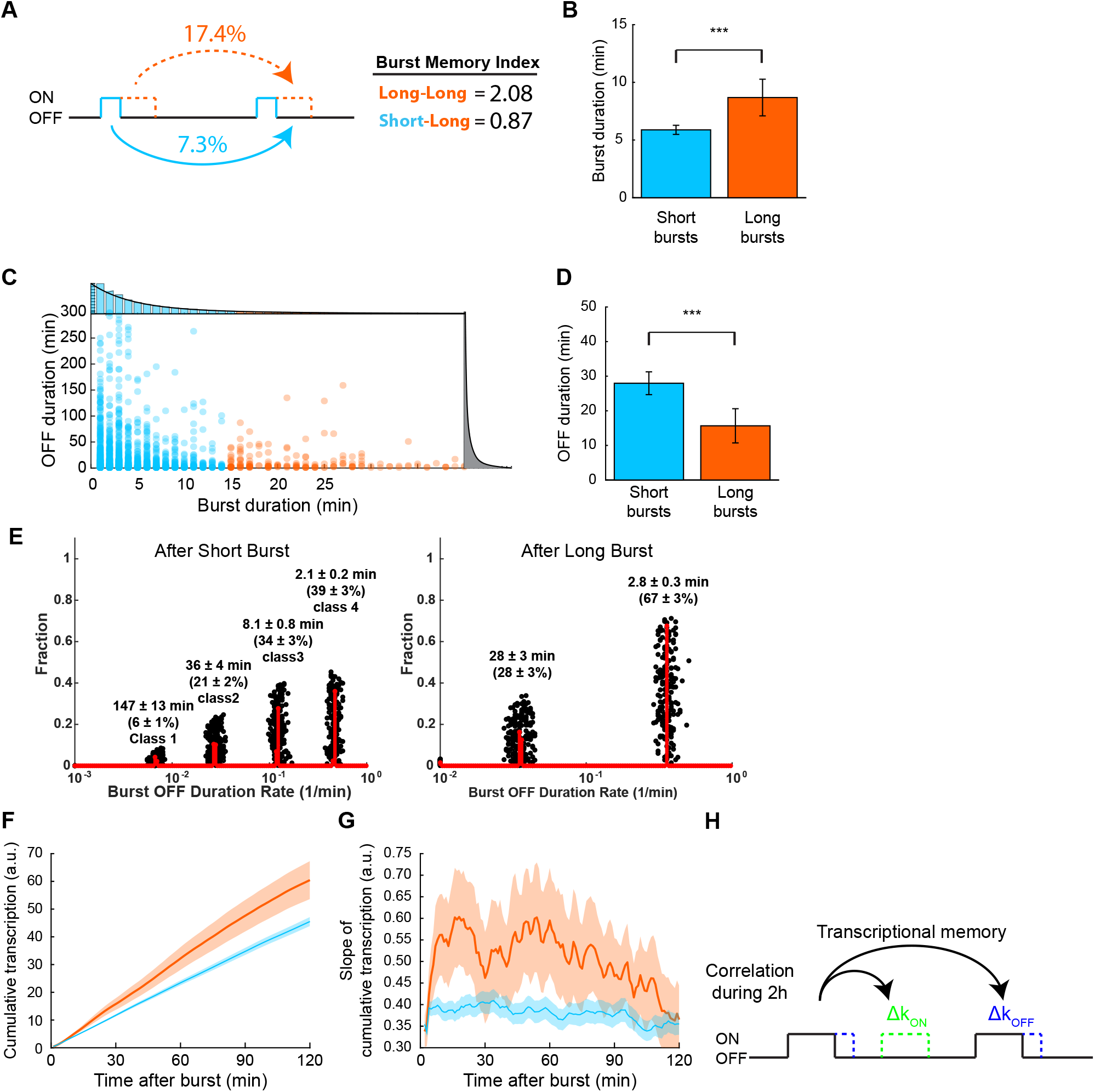
Memory of transcriptional activity stays up to 2 hours. **(A)** A short burst has only 7.3% chance to be followed by a long burst while a long burst has 17.4% chance to be followed by a long burst. **(B)** Quantitation of the ON duration of the subsequent burst after an initial short and long burst. The ON duration of a subsequent burst is significantly longer after a long initial burst compared to a short initial burst. **(C)** Scatter plot of the OFF duration in function of the ON duration. On the top: histogram of the ON duration (1-CDF). On the right: histogram of the OFF duration (1-CDF). The short bursts (15 min or less) are labeled in blue and the long bursts (more than 15 min) are labeled in orange. **(C)** Quantitation of the OFF duration of the subsequent burst after an initial short and long burst. The OFF duration of a subsequent burst is significantly shorter after a long initial burst compared to a short initial burst. **(E)** GRID analysis of the OFF duration immediately after a short (left) and long (right) burst. **(F)** Cumulative transcription of *p21-MS2* after short bursts (blue line) and long bursts (orange line). The average cumulative transcription after long burst is higher than after a short burst (e.g. cells with long bursts produce more mRNA after the long bursts than cells with short bursts). **(H)** Slope of the cumulative transcription. This graph shows the variation of the cumulative slope **(G)** during 2 h. The slope is constant after a short burst but is variable and higher after long bursts up to two hours. **(H)** The burst duration impacts the following bursts duration and frequency for up to 2 hours.

We then took a second approach to quantitatively studying burst duration patterns by examining the average duration of a subsequent burst after a long versus short burst. The average length of a subsequent burst increases nearly 2 fold after a long burst compared with after a short initial burst (**Figure 4B**). Therefore, once a gene is kept on for a period of greater than 15 minutes, the next burst cycle will also likely be long in duration. In contrast, a short burst cycle will less likely lead to a long subsequent transcriptional burst. We suspect that transcription factors and/or the underlying epigenetic marks accumulate at the promoter during the initial long burst cycle, thereby dictating the length of the subsequent burst cycle.

Transcriptional memory is also associated with the frequency of transcription re-initiation. To determine if reinitiation of a burst was dependent on the duration of an initial burst, we plotted the ON duration of a burst versus the subsequent OFF duration. The data indicates that the gene is shut down (OFF) for shorter periods of time after long compared to short bursts (**Figure 4C**). To better quantify differences in re-initiation rates, the average duration of the OFF period following a short versus long burst was calculated. The average duration of the OFF period after a short burst was lengthened nearly 2 fold after a short versus a long burst **(Figure 4D)**. Therefore, a long initial transcriptional burst predominantly supports rapid re-initiation of the subsequent burst cycle. In contrast, a short initial burst cycle is less able to “prime” the gene for subsequent rapid re-initiation of transcription.

To further characterize the OFF periods between two transcriptional bursts, survival curves of the OFF duration after a short versus long burst were analyzed via GRID. There were 4 classes of OFF durations after a short burst and only 2 classes after a long burst (**Figure 4E).** We speculate that these 4 classes are associated with different molecular phenomena related to re-initiation, such as opening of heterochromatin (class 1), local chromatin remodeling/PIC assembly (class 2-3), and RNA Pol II recruitment (class 4). Overall, these results indicate that burst re-initiation is significantly faster after a long burst and likely bypasses select repressive mechanisms (e.g. classes 1 & 3) that are established after a short burst. Transcriptional memory may also be associated with preventing the promoter from entering into a long-lived OFF state likely involving formation of heterochromatin.

Our data suggest that an allele can retain a memory of its own transcriptional activity which will impact its transcriptional kinetics in the future. To measure the duration of this transcriptional memory, we quantified the total mRNA production over time (e.g. cumulative transcription) after short and long bursts (**Figure 4F**). The data indicate that TSs produce significantly more mRNA after a long burst (**Figure 4F**, red line) than after a short burst (**Figure 4F**, blue line). Interestingly, the rate of mRNA production (the slope of the cumulative transcription) is higher for up to two hours after a long burst, suggesting a sustained persistence of the intra-allelic transcriptional memory (**Figure 4G**). Overall, our data show that there is a 2-hour window of transcriptional memory (e.g. Short-Term Transcriptional Memory, STTM) where the bursting duration may affect the future re-initiation (Δk_ON_) and length (Δk_OFF_) of a burst (**Figure 4H**).

### Transcriptional bursting memory can be suppressed by specific stimuli and inhibition of EHMT1/2

To further probe the molecular origins of transcriptional memory, changes in bursting profiles and memory were determined upon treatment of cells with specific stimuli known to activate *p21* gene expression. Previous studies have shown that treatment of cells with Nutlin-3a, UV and RNAi knockdown of EHMT1/2, which dimethylates H3K9 and p53, could enhance expression of *p21(Bunz et al. 1998; Tovar et al. 2006; Chen et al. 2010; Huang et al. 2010*). Therefore, we compared global changes in *p21* MPTB bursting profiles and transcriptional output) using a two state random telegraph model after treatment of cells with Nutlin-3a, UV, and a potent inhibitor of EHMT1/2 (UNC0642, EHMT1/2i)(Liu et al. 2013; Kim et al. 2017). Treatment of cells with Nutlin-3a, UV, and EHMT1/2i globally increased the average burst duration, frequency, and mean transcriptional output from the *p21* gene compared to mock conditions **(Figures 5A-C)**. To determine if elevated p53 levels in the presence of EHMT1/2i could further increase transcriptional output from the *p21* gene, we simultaneously treated the cells with Nutlin-3a and EHMT1/2i. Elevation of p53 levels with Nutlin-3a in the presence of EHMT1/2i increases transcriptional output above levels seen with Nutlin-3a alone **(Figure 5C)**. This is consistent with EHMT1/2’s role in methylating a key residue on p53 (K373) to negatively regulate activation of *p21* transcription(Chen et al. 2010).

**Figure 5.**
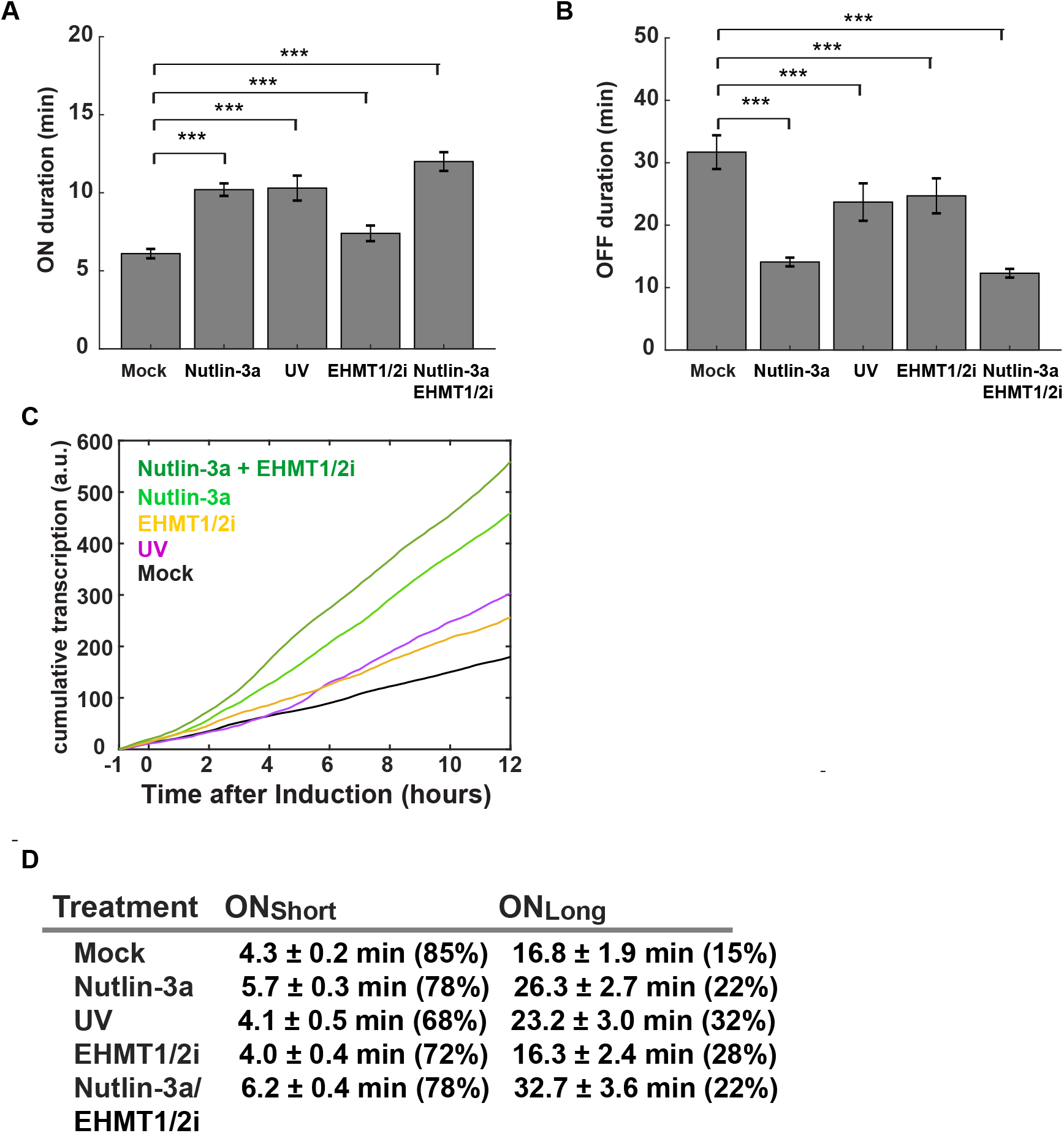
Stimuli specific modulation of bursting profiles. **(A)** ON and **(B)** OFF average duration of bursts based on a two state random telegraph model. **(C)** Cumulative transcription of *p21-MS2*. The average cumulative transcription (above background) show that Nutlin-3a + EHMT1/1i (dark green line, n = 53), Nutlin-3a (light green line, n = 68), UV (magenta line, n = 29), EHMT1/2i (gold line, n = 37) treated cells transcribe the *p21* gene faster than Mock treated cells (black line, n = 85). after Nutlin-3 induction. **(D)** Table showing the percentage and average ON duration from fitting 1-CDF histograms of ON durations using a double exponential decay model from MPTB analysis. p-value *** = < 0.001.

We next compared MPTB ON profiles after the different treatments. In all conditions, treatment of cells with Nutlin-3a, UV, EHMT1/2i and Nutlin-3a + EHMT1/2i, increased the percentage of long bursts **(Figure 5D)**. Only cells treated with Nutlin-3a and Nutlin-3a +EHMT1/2i had a significantly extended duration of short bursts compared to mock treatment. This suggests that MDM2, which is inhibited by Nutlin-3a, may function to restrict the duration of short bursts. All treatments with the exception of EHMT1/2i alone increased the duration of long bursts. This suggests that EHMT1/2 primarily restricts initialization of a long burst at the *p21* gene. This could be due to EHMT1/2 dimethylation of H3K9 or p53 K373. Overall, our results indicate that MPTB bursting profiles can be modulated by a number of different mechanisms.

To determine how the different stimuli impact transcriptional bursting memory, we compared how STTM profiles changed after the various treatments. Remarkably, we see stimuli-specific differences in suppression of bursting memory. The memory index associated with burst durations significantly decreases after UV treatment and EHMT1/2 inhibition compared to basal conditions **(Figure 6A)**. Differences in the ON/OFF durations and transcriptional output after short vs long bursts were also negligible with UV and EHMT1/2 inhibition when compared to basal conditions **(Figures 6B and 6C)**. There were also increases in transcriptional noise across the population of cells after UV treatment and EHMT1/2 inhibition suggesting that STTM and transcriptional noise might be linked **(Figure S4)**. Interestingly, elevation of p53 levels via Nutlin-3a treatment partially reverses the suppression of STTM mediated by EHMT1/2 inhibition **(Figure S6)**. This can be as viewed by increases in the bursting memory index along with restoration of significant differences in ON/OFF durations and transcriptional output after a short versus long burst. Overall, our results suggests that stimuli-specific perturbation of MPTB profiles can suppress the inherent bursting memory regulating basal *p21* expression. Stabilization of p53 via Nutlin-3a is a way to partially restore bursting memory while maximizing transcriptional output at the *p21* gene.

**Figure 6.**
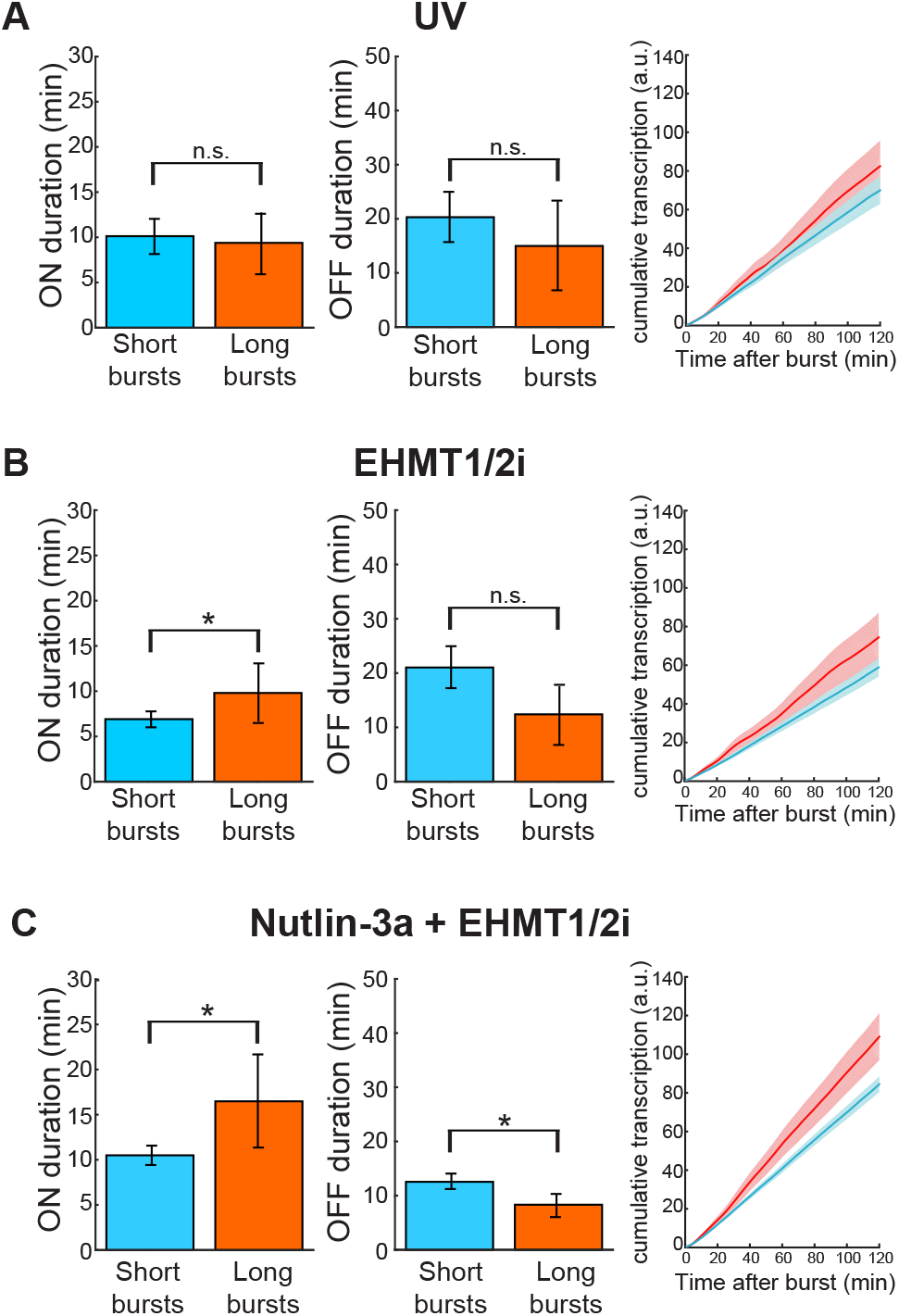
Different Stimuli disrupt STTM profiles. **(A-C)** Quantitation of the ON (left) and OFF (center) durations of the subsequent burst after an initial short and long burst after treating the cells with UV **(A),** EHMT1/2i **(B)**, and Nutlin-3a + EHMT1/2i **(C).** Cumulative transcription of *p21-MS2* (right) after short bursts (blue line) and long bursts (orange line). The average cumulative transcription after long burst is only significantly higher than after a short burst (e.g. cells with long bursts produce more mRNA after the long bursts than cells with short bursts) in cells treated with Nutlin-3a + EHMT1/2i.

## Discussion

We have established an imaging and data analysis pipeline that allows a detailed analysis of dynamic transcriptional bursting patterns at the *p21* cell cycle arrest gene. These analyses provided unique quantitative information on *p21* expression dynamics under basal conditions (e.g. low p53 levels) and after various stimuli that are known to activate *p21* expression (e.g. high p53 levels). Using a random telegraph two state model, we find that elevation of p53 levels increase the probability that cells will harbor a burst of *p21* transcription at any given point in time. This occurs via p53’s ability to increase the burst duration and frequency of bursting at the *p21* gene. Elevated p53 levels were not able to increase transcriptional output above a certain limit, likely due to fundamental limits on the rate of recruitment of Pol II inherent at all genes(Scholes et al. 2017; Rodriguez et al. 2019; Hafner et al. 2020). These results are consistent with a previous live cell imaging study that also studied *p21* bursting dynamics using a similar system(Hafner et al. 2020).

We have extended this previous study by closely examining the ON and OFF durations that respectively correspond to periods of active and inactive *p21* transcription. Multi-exponential fitting methods and GRID analysis allowed us to define multiple ON and OFF states of associated with bursting under basal and activated conditions of *p21* transcription. These multiple ON/OFF states likely reflect the numerous steps associated with turning a gene on and off. For example, going from an OFF to an ON state includes p53 regulated chromatin remodeling to allow access of factors to the enhancer and core promoter regions, PIC formation and repeated Pol II recruitment during convoy formation to extend the burst. These steps are likely interconnected and constantly in flux during a burst, such that disruption of any one step leads to a shut down of the burst. This would be manifested in the appearance of multiple ON states that we see in our data. Indeed, elevated p53 levels can both reduce the percentage of bursts exhibiting a short (~5 min) lifetime while significantly increasing the percentage of events and extending the duration of bursts in the long-lived ON state.

Correspondingly, burst shut down at the *p21* gene also likely occurs via multiple steps. This involves removing histone marks associated with active transcription (e.g. H3K9ac, K14ac and H3K4me3), the subsequent addition of marks required for heterochromatin formation (H3K9me2/3 and H3K27me3) followed by heterochromatin formation(Bannister and Kouzarides 2011). This would be manifested by the appearance of multiple OFF states that we also see in our data. In this manner, elevation of p53 levels are predicted to increase recruitment of histone modifiers that counteract select burst shutdown steps. We also see that elevation of p53 levels eliminates and reduces the prevalence of select OFF states supporting the idea that p53 can act at multiple steps to regulate burst re-initiation or continuation.

Curiously, p53 has also been shown to form a complex with a number of enzymes involved in heterochromatin formation including HDAC1/2, EHMT1, SUV39H1 and LSD1(Juan et al. 2000; Harms and Chen 2007; Huang et al. 2007; Chen et al. 2010). The p53-dependent recruitment of the EHMT1 and SUV39H1 histone methylase complex raises H3K9me2/3 levels at the *p21* promoter(Chen et al. 2010). Therefore, we speculate that p53 can also recruit factors involved in shutting down a burst at the *p21* gene. This suggests that there is a constant balancing act of p53 mediated recruitment of positive and negative regulatory complexes that determine complex bursting patterns and transitions between the multiple ON and OFF states.

By classifying burst events into two distinct classes (e.g. long and short bursts), we could further define the temporal sequence of distinct ON states. We found that select bursting patterns of two sequential long bursts were enriched above random expectation levels in our dataset. Further analysis indicated that future re-initiation rates, burst durations and transcriptional output could be predicted based on the duration of an initial burst, which we now name Short-Term Transcriptional Memory (STTM). This phenomenon is reminiscent of well studied long-term transcriptional memory that is seen over hours and cell generations(Corrigan and Chubb 2014; Cesbron et al. 2015; D’Urso et al. 2016; D’Urso and Brickner 2017; Sood et al. 2017; Palozola et al. 2019). Like the case for long-term transcriptional memory, we speculate that these enriched bursting patterns displaying STTM are likely due to metastable transcription platforms that persist for at least 2 hours. These distinct STTM related platforms are likely associated with unique factors and histone modifications that get rapidly placed but slowly removed as seen in long-term transcriptional memory studies(Corrigan and Chubb 2014; Cesbron et al. 2015; D’Urso et al. 2016; D’Urso and Brickner 2017; Sood et al. 2017; Palozola et al. 2019). In this manner, distinct initiation factors could be rapidly recruited to the STTM platform via these histone marks to give somewhat consistent bursting patterns associated with a defined transcriptional output over a two hour window.

STTM was weakened after UV treatment and inhibition of EHMT1/2 histone methylase activity compared to memory seen during basal expression of the *p21* gene. In cases of cell stress during DNA damage, it is likely that the cell needs to increase production of the *p21* gene as quickly as possible. Our data suggests this may achieved by breaking tightly regulated bursting patterns maintained to keep *p21* levels low in the absence of stress. Elevation of p53 levels with Nutlin-3a in the presence the EHMT1/2 inhibitor partially reverses STTM suppression to levels seen under conditions with Nutlin-3a alone. This suggests that elevation of p53 levels above a threshold not achieved under our UV treatment may serve as a buffer to restore STTM in select cases after it has become dysregulated.

Interestingly, transcriptional noise across the population of cells also temporally increased after UV treatment and inhibition of EHMT1/2 histone methylase activity. The fact that STTM is the strongest and transcriptional noise the lowest under basal conditions may suggest that the expression of *p21* at low p53 levels needs to be tightly regulated. Basal expression of the p21 protein is thought to perform a balancing act by regulating populations of cycling and quiescent cells(Overton et al. 2014). This tight regulation of basal *p21* expression via STTM may explain how p21 functions in stem cells to maintain the balance between dormant and self-renewing cells in niches. In such a scenario, extreme variations in STTM and transcriptional noise could stochastically switch a cycling stem cell into a dormant one and vice versa via select cues or stimuli. This process of cell phenotype switching could be initiated by similar mechanisms as seen when this balance is broken during DNA damage after p53 induces *p21* expression to initiate cell cycle arrest. Cancer also finds a way to disrupt this balancing act by inactivating p53’s activity via multiple mechanisms to suppress *p21* expression leading to uncontrollable cycling of cells and tumor growth. So it will be interesting to see how STTM at the *p21* gene is affected in cancer models. We next aim to use our data analysis methods to examine bursting patterns of additional genes in human and yeast cells to determine if bursting memory is an inherent feature associated with tight regulation of gene expression.

## MATERIAL AND METHODS

### Cell culture

Human U2OS cells (American Type Culture Collection [ATCC], HTB-96) were grown at 37°C and 5% CO2 in Dulbecco’s Modified Eagle Medium (DMEM) high glucose (Corning) containing 10% Fetal Bovine Serum (FBS - Atlanta Biologicals), and 1% penicillin–streptomycin (Gibco), according to ATCC recommendations.

### Plasmids, genome editing, and viral transduction

We created a U2OS cell line containing 24 MS2-SL into the 3’UTR of the endogenous p21 gene and expressing MCP-GFP explained in detail within our previous study (Carvajal et al., 2018).

To over-express p53wt and p53-R273H, we introduced p53wt-HALO and p53R273H-HALO into Tet-On 3G inducible expression retroviral systems. We use PCR to amplify the p53wt and p53-R273H sequences using the following primers (ATCGGGATCCTATGGAGGAGCCGCAGTCAG and ATCGGAATTCGCCCTTTCACTAAGTGGTTG) from CMV driven expression plasmids containing the sequences for human p53 WT (pP53WT-Halo) and a p53 mutant (pP53R273H-Halo) that were c-terminally tagged with the HaloTag protein sequence (details of plasmids are available upon request). The plasmid pRetroX-Tight-Pur (Clontech) and the PCR products have been digested with BamHI and EcoRI, the plasmid has been dephosphorylated with FastAP Thermosensitive Alkaline Phosphatase (Thermofisher) and they have been ligated together with T4 DNA ligase (NEB). Either the plasmids pRetro-TRE3G-p53wt-HALO-Puro or pRetro-TRE3G-p53R270H-HALO-Puro have been co-transfected with pQEXIN-tetON3G following the Clonetech retroviral user manual into 13A cells. After 10μM of Doxycycline induction, the 13A-p53wt and 13A-p53R273H start to express the p53 proteins.

The plasmids pUC19-p21-v5-MS2-Neo-p21, pRetro-TRE3G-p53wt-HALO-Puro and pRetro-TRE3G-p53R270H-HALO-Puro are available on Addgene.

### Live Cell Imaging

For live cell imaging, cells were washed in PBS and were imaged at 37 °C in Leibovitz’s L-15 Medium (Gibco) containing 10% FBS. Time-lapse z-series images were captured on a wide-field microscope (Olympus IX-81 stand) equipped with an electron multiplying CCD camera (iXon3 DU-897E-CS0-#BV; Andor) and controlled by MetaMorph software (Molecular Devices, Sunnyvale, CA). A 491-nm laser (Calypso-25; Cobolt) was used for illumination. The microscope also was equipped with an automated XY stage (MS2000-XY with an extra-fine lead-screw pitch of 0.635 mm and a 10-nm linear encoder resolution; Applied Scientific Instrumentation) and a piezo-Z stage (Applied Scientific Instrumentation) for fast z-stack acquisition. The cells were kept at 37 °C with a stage top incubator (INUBH-ZILCS-F1; Tokai Hit). An objective with a 1.4 NA and 60X magnification yielded a pixel-size of 266.8nm. Cells expressing the MCP-GFP were imaged for 15 hours with 11 stacks of 500 nm (1 picture every minute). Media was replaced under the microscope after 3 hours with fresh L15 media containing 10% FBS with a mock solution (1/1000 of DMSO - Sigma D8418), 10μM Nutlin-3 (Sigma N6287), 1μg/mL Doxycycline (Sigma D9891) or 500nM UNC0642 (Sigma SML1037). For UV treatment, cells were pulsed for 50ms every minute with 0.5W/cm^2^ from a 405-nm laser.

Images were analyzed after a maximum intensity projection along Z in ImageJ (http://rsb.info.nih.gov/ij/). The TS fluorescence was quantitated with our custom MATLAB software (STAR-Burst). The TS of each cell was detected semi-automatically and the fluorescence signal measured for each time point. Fast fluctuations in the fluorescence signal of the TS were removed using a rolling average of 3 minutes. Transcriptional activity was normalized by dividing the fluorescence of the TS by the background signal. The TS was deemed active when transcriptional activity was higher than 1.5 fold above the background.

### Statistical analysis

#### p-values and confidence interval

The p-value have been calculated using the Student’s t-test. The confidence intervals have been calculated using the Student t distribution for 2.5% and 97.5%.

### Plots of Cumulative transcription and Transcription Noise

We subtracted 1 to the fold change to center the fluorescence intensity of each TS to 0. We then calculated the accumulated TS intensity over time for each cell and plotted the average rate of cumulative transcription within the population of cells over time. Transcriptional noise was calculated by determining the mean and standard deviation of fluorescent intensity of the TS at each time point during 15 hours of imaging. The coefficient of variation squared (CV^2^) was then calculated by taking the square of the standard deviation divided by the square of the mean for each time point and plotting the average CV^2^ over a sliding 60 minute window as a function of time.

### Colormap

We normalized the fluorescence intensity of each condition with 0 for the lower intensity quantified and 1 for the highest. We converted each value to an RGB value (colormap fire – see legend of Figure 1E).

### ON probability for the two state random telegraph model

We quantify the proportion of cells containing a burst at each time point with the data averaged over a 2-hour sliding window (1 hour before the time point and 1 hour after).

### Burst ON duration and OFF duration for the two state random telegraph model

For each condition (control, Nutlin-3a and p53wt overexpression), we calculated the average burst duration and delay between bursts at each time point over a 2-hour window starting 2 hours after treating with Nutlin-3a or doxocycline to initiate overexpression of p53.

### Bootstrapping

We performed a bootstrapping resampling analysis(Haimovich et al. 2013) to determine the 95% confidence interval of ON probability, burst duration, and OFF duration. Each time, we selected 10 000 groups of transcription sites containing the same number of the original data. We calculated the average value for each groups and plotted the 95% confidence interval as gray or colored area.

### Burst duration and OFF duration over time

We calculated, for each conditions, the average burst duration and delay between bursts for a 4-hour windows (2 hours before the time point and 2 hours after).

### Multi-exponential fitting and GRID analysis of ON and OFF durations

1-CDF histograms of ON durations accumulated from a number of cells (listed in each figure legend) were fitted to a single and double exponential decay model. Error values for the percentage of events in each class and the average duration of each class were based on the standard error of the mean. Comparison of chi-squared values assessing the variance of the exponential fit from the histogram data indicated that a double exponential decay model was a better fit to the data in all cases. GRID analysis of the ON and OFF durations were performed on survival curves as described in(Reisser et al. 2020) and confirmed results from the multi-exponential fitting. Resampling of 80% of the data was performed 100 times to generate error values for the percentage of events in each class and the average duration of each class.

### Classification of burst events into short and long bursts

Multi-exponential fitting with a double exponential decay model was used to determine the percentage and average duration of ON bursts in the two populations of each condition (control, Nutlin-3a, UV, EHMT1/2 inhibition and Nutlin-3a + EHMT1/2 inhibition). Using the percentage of events (A_S_) and average duration (kS) of the population of events with the shortest average burst duration, we used a single exponential decay model (y=A_S_*exp^-kS*t^) to calculate the time threshold at which only 2.5% of shortest class of bursts were seen in the 1-CDF histogram. Events less than or equal to this time threshold were classified as short bursts while events longer than this threshold were classified as long bursts.

### Calculation of transcriptional memory profiles

To determine the burst memory index, we first calculated the observed percentage of long burst ON events that were subsequent to either a short versus long initial burst in a dataset for each treatment condition. We then divided the observed value for a long-long and short-long series of bursts by the expected percentage of total burst events in the longest class of burst durations as determined by multi-exponential fitting of 1-CDF histograms of our ON burst durations. We plotted the average cumulative transcription rate within a 2 hour window after each short or long burst event in all datasets for each condition with the 97.5% confidence intervals determined by a student’s t-test. To determine the time of persistence of memory, the slope of the cumulative transcription rate over subsequent time points was plotted over a 2 hour time window with the 97.5% confidence intervals determined by a student’s t-test.

## Supporting information

Video of p21MS2 with Nutlin-3a

## Data and software availability

Code for STAR-burst is available at //github.com/Coleman-Laboratory/STAR-burst.

## ACKNOWLEDGMENTS

We would like to acknowledge the current and past members of the Coleman and Singer labs for reagents, technical assistance and critically reviewing this manuscript. Funding was obtained as part of the 4D Nucleome via NIH/NIDA 8U01DA047729 (RHS and RAC), NIH NS 083085 (RHS), NIH/NIGMS 1R35GM136296 (RHS), and NIH/NIGMS 1R01GM126045 (RAC).

## AUTHOR CONTRIBUTIONS

AS, RHS and RAC conceived the overall project and experimental design. AS generated the cell lines and performed the experiments. AS designed the code for STAR-burst. AS and RAC performed data analysis. AS, RHS and RAC wrote the manuscript. RAC and RHS obtained funding for the studies.

---

**Figure S1.**
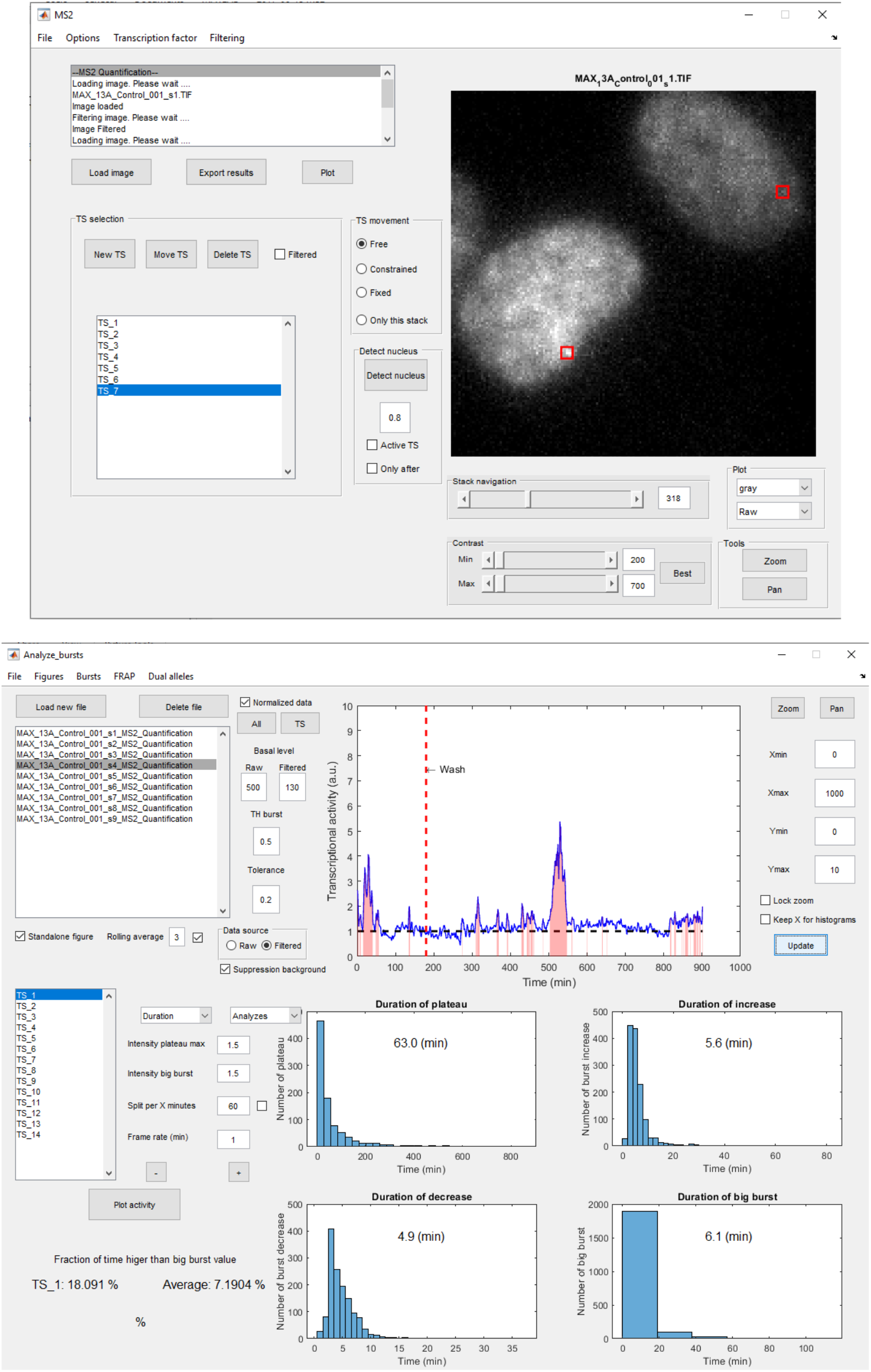
MS2-Quant and Analyze burst – tools for analyzing MS2 transcriptional activity in MATLAB.

**Figure S2.**
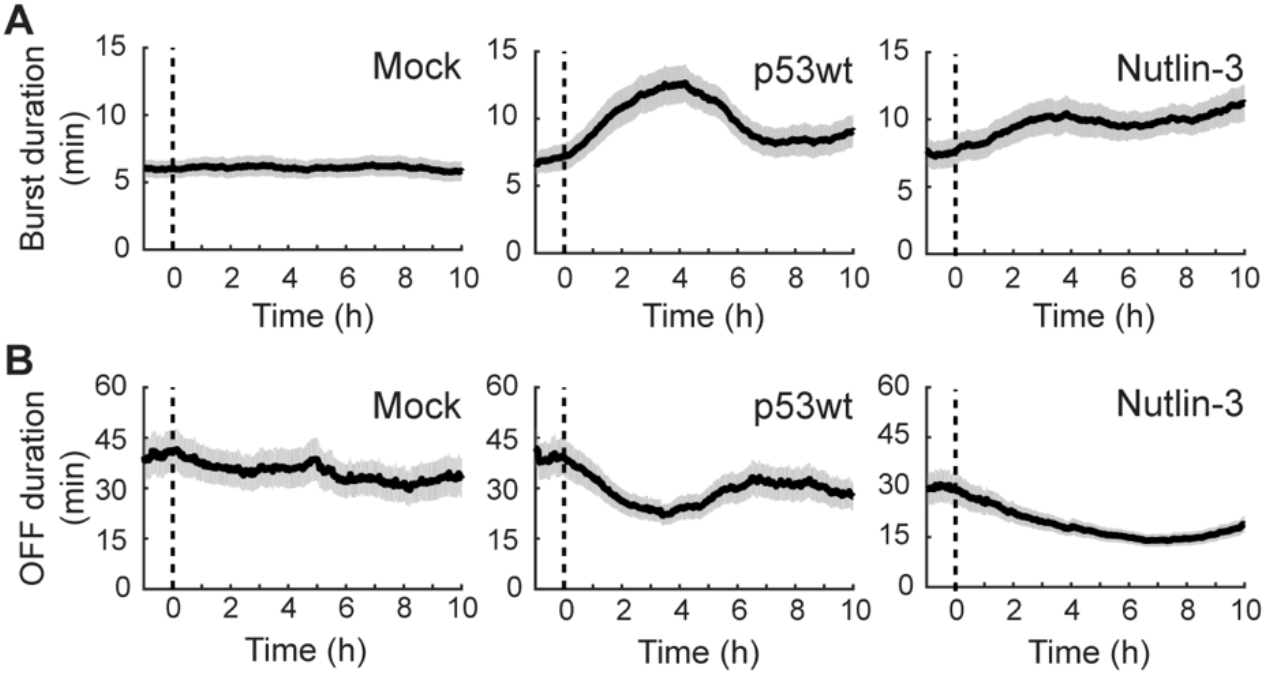
p53 increase p21 expression by enhancing burst duration and frequency. **(A and B)** Average Burst **(A)** and OFF **(B)** duration over-time. The Burst and OFF duration is calculated for a 4h windows (2 hours before and 2 hours after each time point). At 0h, the cells are treated with a Mock (DMSO), Doxycycline or Nutlin-3a.

**Figure S3.**
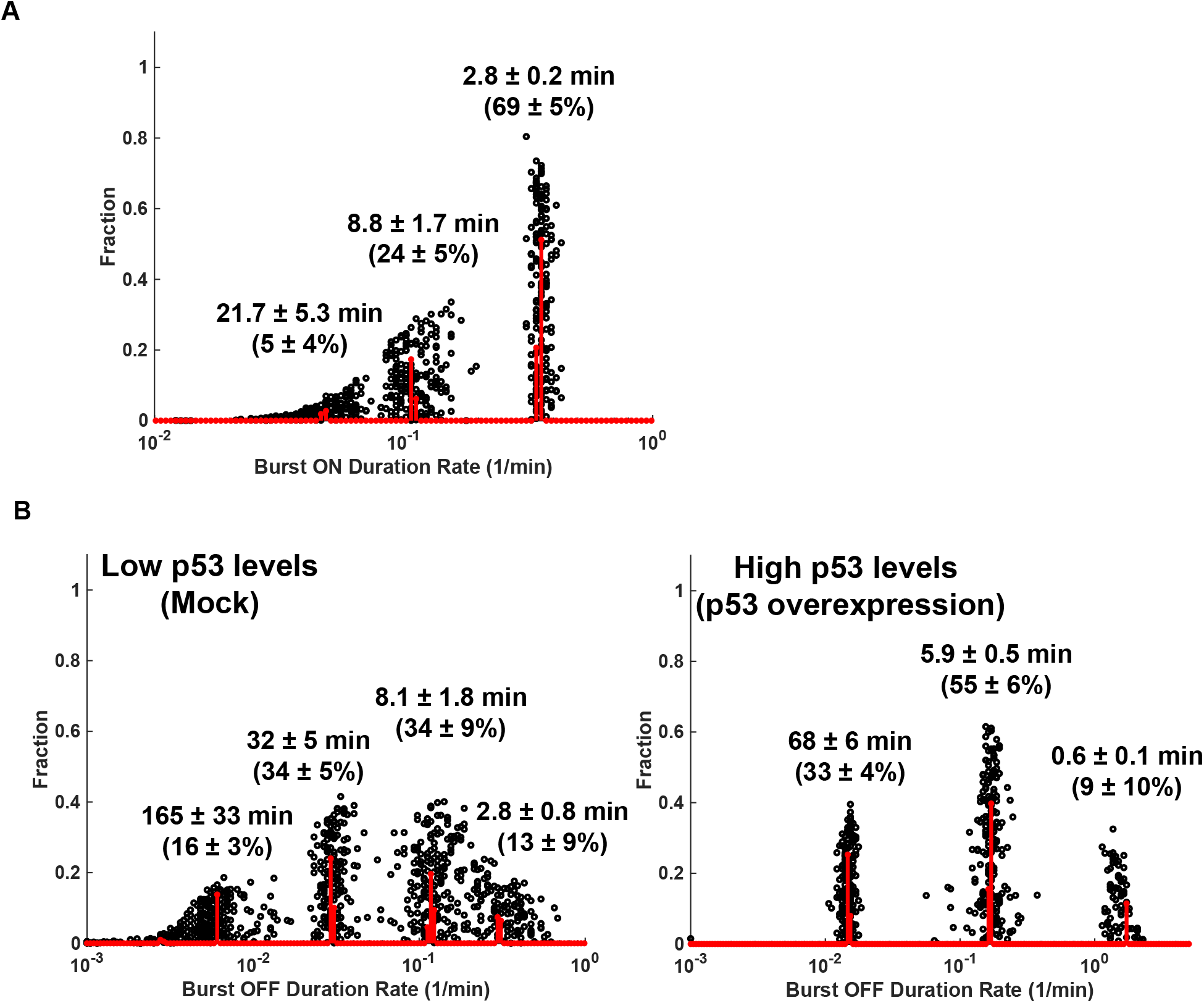
GRID analysis indicates multiple ON and OFF states. **(A)** GRID analysis of the ON duration of bursts in Mock treated cells. Values for average duration shown have been converted by inverting the Burst ON duration rate of each population **(B)**. GRID analysis of the OFF duration of bursts in Mock treated cells (left) and p53 overexpressing cells (right). Values for average duration shown have been converted by inverting the Burst OFF duration rate of each population.

**Figure S4.**
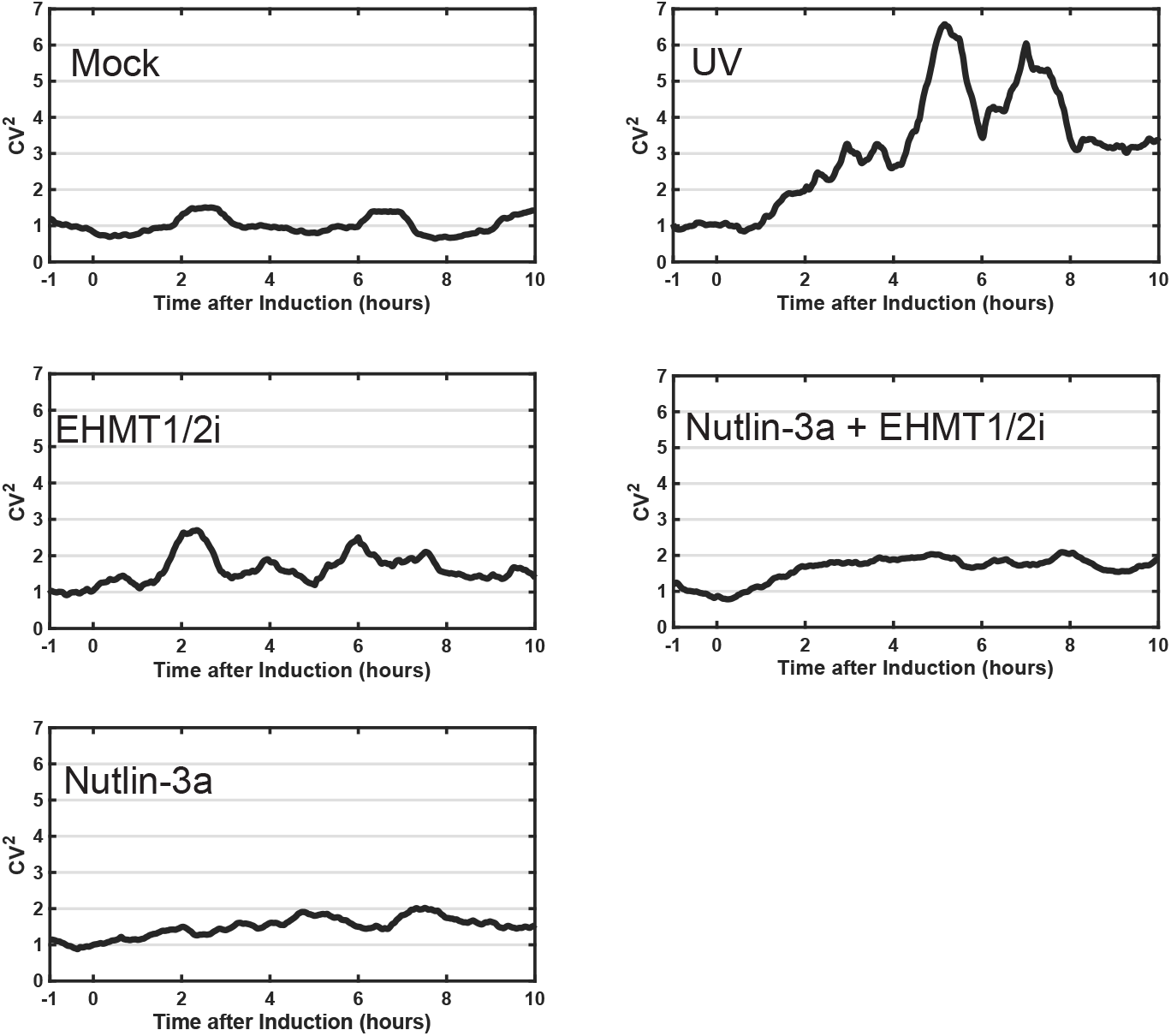
Transcriptional Noise increases after various stimuli. Quantitation of the square of the Coefficient of Variance (σ^2^/mean^2^, CV^2^) of transcription spot fluorescence intensity as a function of time across cells in a population. The CV^2^ was calculated based on the average from a sliding 60 minute window and normalized to 1 based on the mean CV^2^ from 1 hour prior to induction with the various stimuli. Mock (85 cells), UV (29 cells), EHMT1/2i (37 cells), Nutlin-3a + EHMT1/1i (53 cells), Nutlin-3a (68 cells).

**Figure S5.**
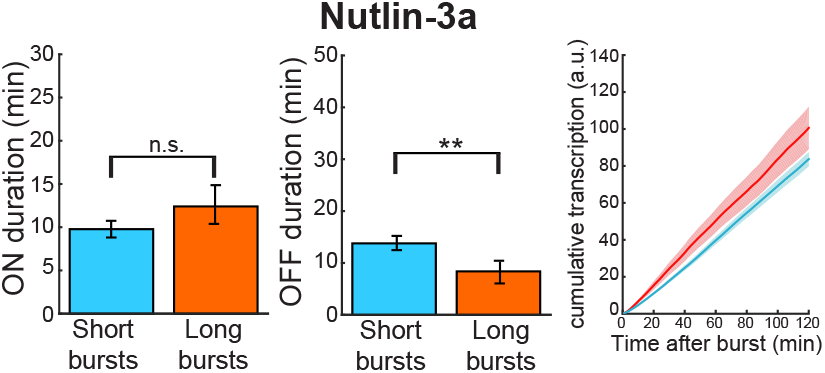
Nutlin-3a weakly suppresses STTM. Quantitation of the ON (left) and OFF (center) durations of the subsequent burst after an initial short and long burst after treating the cells with Nutlin-3a. Cumulative transcription of *p21-MS2* (right) after short bursts (blue line) and long bursts (orange line). The average cumulative transcription after long burst is only slightly significantly higher than after a short burst (e.g. cells with long bursts produce more mRNA after the long bursts than cells with short bursts).

## REFERENCES

Bannister AJ, Kouzarides T. 2011. Regulation of chromatin by histone modifications. Cell Res 21: 381–395.

Biswas J, Li W, Singer RH, Coleman RA. 2021. Imaging Organization of RNA Processing within the Nucleus. Cold Spring Harbor perspectives in biology.

Bullock AN, Henckel J, Fersht AR. 2000. Quantitative analysis of residual folding and DNA binding in mutant p53 core domain: definition of mutant states for rescue in cancer therapy. Oncogene 19: 1245–1256.

Bunz F, Dutriaux A, Lengauer C, Waldman T, Zhou S, Brown JP, Sedivy JM, Kinzler KW, Vogelstein B. 1998. Requirement for p53 and p21 to sustain G2 arrest after DNA damage. Science (New York, NY) 282: 1497–1501.

Carvajal LA, Neriah DB, Senecal A, Benard L, Thiruthuvanathan V, Yatsenko T, Narayanagari SR, Wheat JC, Todorova TI, Mitchell K et al. 2018. Dual inhibition of MDMX and MDM2 as a therapeutic strategy in leukemia. Sci Transl Med 10.

Cesbron F, Oehler M, Ha N, Sancar G, Brunner M. 2015. Transcriptional refractoriness is dependent on core promoter architecture. Nature communications 6: 6753.

Chang GS, Chen XA, Park B, Rhee HS, Li P, Han KH, Mishra T, Chan-Salis KY, Li Y, Hardison RC et al. 2014. A comprehensive and high-resolution genome-wide response of p53 to stress. Cell reports 8: 514–527.

Chen L, Li Z, Zwolinska AK, Smith MA, Cross B, Koomen J, Yuan ZM, Jenuwein T, Marine JC, Wright KL et al. 2010. MDM2 recruitment of lysine methyltransferases regulates p53 transcriptional output. The EMBO journal 29: 2538–2552.

Corrigan AM, Chubb JR. 2014. Regulation of transcriptional bursting by a naturally oscillating signal. Current biology: CB 24: 205–211.

Corrigan AM, Tunnacliffe E, Cannon D, Chubb JR. 2016. A continuum model of transcriptional bursting. eLife 5.

D’Urso A, Brickner JH. 2017. Epigenetic transcriptional memory. Curr Genet 63: 435–439.

D’Urso A, Takahashi YH, Xiong B, Marone J, Coukos R, Randise-Hinchliff C, Wang JP, Shilatifard A, Brickner JH. 2016. Set1/COMPASS and Mediator are repurposed to promote epigenetic transcriptional memory. eLife 5.

de Jonge WJ, O’Duibhir E, Lijnzaad P, van Leenen D, Groot Koerkamp MJ, Kemmeren P, Holstege FC. 2017. Molecular mechanisms that distinguish TFIID housekeeping from regulatable SAGA promoters. The EMBO journal 36: 274–290.

Donczew R, Warfield L, Pacheco D, Erijman A, Hahn S. 2020. Two roles for the yeast transcription coactivator SAGA and a set of genes redundantly regulated by TFIID and SAGA. eLife 9.

Donner AJ, Hoover JM, Szostek SA, Espinosa JM. 2007. Stimulus-specific transcriptional regulation within the p53 network. Cell Cycle 6: 2594–2598.

Eldar A, Elowitz MB. 2010. Functional roles for noise in genetic circuits. Nature 467: 167–173.

Hafner A, Reyes J, Stewart-Ornstein J, Tsabar M, Jambhekar A, Lahav G. 2020. Quantifying the Central Dogma in the p53 Pathway in Live Single Cells. Cell Syst 10: 495–505 e494.

Haimovich G, Medina DA, Causse SZ, Garber M, Millan-Zambrano G, Barkai O, Chavez S, Perez-Ortin JE, Darzacq X, Choder M. 2013. Gene expression is circular: factors for mRNA degradation also foster mRNA synthesis. Cell 153: 1000–1011.

Harms KL, Chen X. 2007. Histone deacetylase 2 modulates p53 transcriptional activities through regulation of p53-DNA binding activity. Cancer research 67: 3145–3152.

Huang J, Dorsey J, Chuikov S, Zhang X, Jenuwein T, Reinberg D, Berger SL. 2010. G9a and Glp methylate lysine 373 in the tumor suppressor p53. The Journal of biological chemistry 285: 9636–9641.

Huang J, Sengupta R, Espejo AB, Lee MG, Dorsey JA, Richter M, Opravil S, Shiekhattar R, Bedford MT, Jenuwein T et al. 2007. p53 is regulated by the lysine demethylase LSD1. Nature 449: 105–108.

Huisinga KL, Pugh BF. 2004. A genome-wide housekeeping role for TFIID and a highly regulated stress-related role for SAGA in Saccharomyces cerevisiae. Molecular cell 13: 573–585.

Juan LJ, Shia WJ, Chen MH, Yang WM, Seto E, Lin YS, Wu CW. 2000. Histone deacetylases specifically down-regulate p53-dependent gene activation. The Journal of biological chemistry 275: 20436–20443.

Kim Y, Lee HM, Xiong Y, Sciaky N, Hulbert SW, Cao X, Everitt JI, Jin J, Roth BL, Jiang YH. 2017. Targeting the histone methyltransferase G9a activates imprinted genes and improves survival of a mouse model of Prader-Willi syndrome. Nat Med 23: 213–222.

Kubik S, Bruzzone MJ, Shore D. 2017. TFIID or not TFIID, a continuing transcriptional SAGA. The EMBO journal 36: 248–249.

Larsson AJM, Johnsson P, Hagemann-Jensen M, Hartmanis L, Faridani OR, Reinius B, Segerstolpe A, Rivera CM, Ren B, Sandberg R. 2019. Genomic encoding of transcriptional burst kinetics. Nature 565: 251–254.

Lenstra TL, Rodriguez J, Chen H, Larson DR. 2016. Transcription Dynamics in Living Cells. Annual review of biophysics 45: 25–47.

Lev Bar-Or R, Maya R, Segel LA, Alon U, Levine AJ, Oren M. 2000. Generation of oscillations by the p53-Mdm2 feedback loop: a theoretical and experimental study. Proceedings of the National Academy of Sciences of the United States of America 97: 11250–11255.

Liu F, Barsyte-Lovejoy D, Li F, Xiong Y, Korboukh V, Huang XP, Allali-Hassani A, Janzen WP, Roth BL, Frye SV et al. 2013. Discovery of an in vivo chemical probe of the lysine methyltransferases G9a and GLP. Journal of medicinal chemistry 56: 8931–8942.

Molina N, Suter DM, Cannavo R, Zoller B, Gotic I, Naef F. 2013. Stimulus-induced modulation of transcriptional bursting in a single mammalian gene. Proceedings of the National Academy of Sciences of the United States of America 110: 20563–20568.

Overton KW, Spencer SL, Noderer WL, Meyer T, Wang CL. 2014. Basal p21 controls population heterogeneity in cycling and quiescent cell cycle states. Proceedings of the National Academy of Sciences of the United States of America 111: E4386–4393.

Palozola KC, Lerner J, Zaret KS. 2019. A changing paradigm of transcriptional memory propagation through mitosis. Nature reviews Molecular cell biology 20: 55–64.

Patange S, Girvan M, Larson DR. 2018. Single-cell systems biology: probing the basic unit of information flow. Curr Opin Syst Biol 8: 7–15.

Raj A, van Oudenaarden A. 2008. Nature, nurture, or chance: stochastic gene expression and its consequences. Cell 135: 216–226.

Reisser M, Hettich J, Kuhn T, Popp AP, Grosse-Berkenbusch A, Gebhardt JCM. 2020. Inferring quantity and qualities of superimposed reaction rates from single molecule survival time distributions. Scientific reports 10: 1758.

Rodriguez J, Larson DR. 2020. Transcription in Living Cells: Molecular Mechanisms of Bursting. Annu Rev Biochem 89: 189–212.

Rodriguez J, Ren G, Day CR, Zhao K, Chow CC, Larson DR. 2019. Intrinsic Dynamics of a Human Gene Reveal the Basis of Expression Heterogeneity. Cell 176: 213–226 e218.

Scholes C, DePace AH, Sanchez A. 2017. Combinatorial Gene Regulation through Kinetic Control of the Transcription Cycle. Cell Syst 4: 97–108 e109.

Senecal A, Munsky B, Proux F, Ly N, Braye FE, Zimmer C, Mueller F, Darzacq X. 2014. Transcription factors modulate c-Fos transcriptional bursts. Cell reports 8: 75–83.

Sood V, Cajigas I, D’Urso A, Light WH, Brickner JH. 2017. Epigenetic Transcriptional Memory of GAL Genes Depends on Growth in Glucose and the Tup1 Transcription Factor in Saccharomyces cerevisiae. Genetics 206: 1895–1907.

Timmers HTM. 2021. SAGA and TFIID: Friends of TBP drifting apart. Biochim Biophys Acta Gene Regul Mech 1864: 194604.

Tovar C, Rosinski J, Filipovic Z, Higgins B, Kolinsky K, Hilton H, Zhao X, Vu BT, Qing W, Packman K et al. 2006. Small-molecule MDM2 antagonists reveal aberrant p53 signaling in cancer: implications for therapy. Proceedings of the National Academy of Sciences of the United States of America 103: 1888–1893.

Tunnacliffe E, Chubb JR. 2020. What Is a Transcriptional Burst? Trends Genet 36: 288–297.

Vassilev LT, Vu BT, Graves B, Carvajal D, Podlaski F, Filipovic Z, Kong N, Kammlott U, Lukacs C, Klein C et al. 2004. In vivo activation of the p53 pathway by small-molecule antagonists of MDM2. Science (New York, NY) 303: 844–848.

Willis A, Jung EJ, Wakefield T, Chen X. 2004. Mutant p53 exerts a dominant negative effect by preventing wildtype p53 from binding to the promoter of its target genes. Oncogene 23: 2330–2338.

